# Field-derived temperature correction compromises eDNA-based abundance inference

**DOI:** 10.64898/2026.07.03.735744

**Authors:** Martin Ogonowski, Zandra Gerdes

## Abstract

Environmental DNA (eDNA) has emerged as a promising tool for estimating fish abundance, yet linking eDNA concentration to true density remains a significant challenge in seasonal systems, where the signal is strongly influenced by temperature. We investigated whether eDNA can serve as an abundance index for three-spined stickleback (*Gasterosteus aculeatus*) in four coastal bays of the Baltic Sea (5.7–20.5°C, April–July 2023), by pairing eDNA sampling with two trap types of contrasting catchability. Light traps capture fish by phototactic attraction during darkness, so their catchability is driven primarily by night duration rather than temperature, while benthic traps respond to temperature through the same activity-driven mechanism as eDNA production. The temperature sensitivity of eDNA estimated from field data was far higher than physiological expectation (Q10 = 12.4, against a maximum metabolic rate benchmark of Q10 = 3.5), indicating that the field temperature signal reflects ecological change in addition to metabolism. We then compared how well three eDNA predictors tracked a combined trap-based abundance index: uncorrected eDNA, eDNA corrected with the temperature response constrained to the laboratory metabolic rate (a first-principles correction), and eDNA corrected with the response estimated from the field data. Uncorrected and first-principles-corrected eDNA were both strong predictors of abundance (standardised slopes of 0.45 and 0.43), whereas the field-corrected predictor was not (0.08). Uncorrected and first-principles-corrected eDNA performed comparably because temperature and abundance increased together over the season; the first-principles correction is nonetheless preferable, as it remains reliable when this covariation is unknown a priori. We conclude that estimating a temperature correction from field data should be avoided in seasonal eDNA monitoring, because it removes the abundance signal together with the temperature effect and assumes a stability in abundance that cannot be verified without independent reference data.

## 1. Introduction

Monitoring fish populations is essential for effective management, yet no single sampling method provides an unbiased estimate of abundance. In shallow coastal ecosystems, passive gillnets and benthic traps are the most common tools for tracking littoral fish, but neither is fully quantitative: both are size-selective, and their catchability depends strongly on the physical activity of the target species, which in ectotherms is governed by water temperature and species-specific behaviour (Linløkken and Haugen 2006, Ward et al. 2012).

Environmental DNA (eDNA), the genetic material shed by organisms into their surroundings, has transformed aquatic biodiversity monitoring by enabling species detection without physical capture (Ficetola et al. 2008, Jerde et al. 2011, Langlois et al. 2021). As the methodology has matured, attention has shifted toward the more ambitious goal of estimating abundance from eDNA concentration, typically via quantitative PCR (qPCR; Yates et al. 2019, Spear et al. 2021, Pont et al. 2023). This prospect is especially attractive in aquatic systems, where conventional active sampling is logistically difficult, environmentally damaging, or heavily gear-selective.

The relationship between eDNA concentration and true fish abundance is not straightforward, however. The amount of DNA detectable in a water sample depends partly on the biomass of fish but also on how much DNA is released per unit biomass. Because fish are ectotherms: their metabolism scales with temperature, so the same individual sheds more DNA in warm water simply because it is more metabolically active (Jo et al. 2019a, Thalinger et al. 2021, Jo 2025). At the same time, DNA degradation accelerates with warming as microbial activity intensifies (Tsuji et al. 2017, Jo et al. 2019b, McCartin et al. 2022). Moreover, in open coastal, tidal or riverine systems, water exchange continuously removes eDNA before local shedding and degradation rates can balance. Measured eDNA therefore reflects primarily how fast fish are producing it at that moment (Sassoubre et al. 2016, Jo et al. 2019b), a rate that rises steeply with water temperature. This amplifies the temperature signal in the data and makes it harder to separate a genuine change in fish numbers/biomass from a temperature-driven change in shedding rate, especially when sampling spans a wide seasonal temperature range.

This thermal sensitivity of eDNA production has been grounded in the Metabolic Theory of Ecology (MTE; Brown 2004), which predicts that biological rates scale with temperature according to allometric principles. Yates et al. (2025) recently formalized the relationship, showing that eDNA production can be modelled as a function of abundance, allometrically scaled body mass, and a temperature dependency function f(T) analogous to the consumption functions used in bioenergetics models (Stewart and Ibarra 1991, Jobling 1994, Leloutre et al. 2018). They noted that no study to date has examined eDNA production across a temperature gradient wide enough to characterise the shape of f(T) but proposed that field data spanning a sufficient natural gradient could, in principle, provide this characterisation. Any temperature correction estimated from seasonal field data conflates two inseparable signals: the true effect of temperature on the specific eDNA production, and the co-occurring change in abundance, since both track the same seasonal temperature gradient. Disentangling these components demands an independent abundance reference that is itself free of temperature bias. Existing laboratory experiments have typically examined only two or three temperature points (Klymus et al. 2015, Lacoursière-Roussel et al. 2016, Jo et al. 2019b, Hervé et al. 2023), too few to characterise the functional form of the temperature – eDNA relationship across ecologically relevant ranges.

A further complication, largely unaddressed in the eDNA literature, is that the conventional trap-based methods used to validate eDNA are themselves subject to temperature-dependent catchability biases. Passive gears that rely on fish actively encountering them like benthic fyke nets or gillnets, for example, systematically underestimate abundance when water is cold and fish are inactive, even where those fish are present at high density (Arreguín-Sánchez 1996, Ward et al. 2012). Ward et al. (2012) demonstrated a 7-fold variation in gillnet catchability across a ∼7°C window when abundance was held approximately constant by sampling at a fixed phenological stage, confirming that temperature-driven catchability variation in passive gears must be accounted for. This creates a circularity for eDNA validation: if both the eDNA signal and the conventional abundance index are amplified by temperature through the same metabolic–activity mechanism, comparing them inflates the apparent eDNA–abundance relationship without revealing the thermal bias in either method.

While active gears also have been used as eDNA references, including bottom-trawls (Salter et al. 2019, Maes et al. 2024) and standardized angling for northern pike (Ogonowski et al. 2023), they are not immune to the same confound. Ogonowski et al. (2023) found the eDNA–angling relationship for pike was temperature-dependent, significant only at higher temperatures coinciding with spawning. With a single reference, however, it is impossible to determine whether the eDNA signal, the reference catchability, or both were responding to temperature. To our knowledge, no study has combined eDNA with reference gears chosen to have contrasting responses to temperature, a design that allows the thermal component of the eDNA signal to be separated from the abundance signal rather than left unresolved.

In this study, we exploit the opposing temperature sensitivities of two trap types to characterise the observation process underlying the eDNA signal and to test whether usable abundance information can be recovered through physiologically constrained correction.

Working with three-spined stickleback (*Gasterosteus aculeatus*) in four coastal brackish bays of the Baltic Sea across a full spring-to-summer season (5.7–20.5°C), we paired quantitative eDNA sampling with light traps and benthic traps deployed concurrently. Light traps capture fish via phototaxis at night, so their catchability is governed primarily by night duration rather than water temperature, making them the least temperature-confounded abundance reference available in this system (Ogonowski 2026). Benthic traps capture fish passively, so their catchability increases with temperature through the same metabolic-activity pathway that drives eDNA shedding. By standardising each trap type for its primary bias — night duration for light traps and temperature for benthic traps — we derived two corrected abundance indices operating on orthogonal seasonal axes.

This design allowed us to address two questions. First, can the opposing catch patterns of a thermally driven and a non-thermally driven method resolve the temperature component of the eDNA signal? We hypothesized that if the temperature effect on eDNA is substantial, raw eDNA concentration should correlate positively with the temperature-dependent method (benthic traps) and negatively with the temperature-independent method (light traps). Second, we asked whether the field-estimated temperature dependence is consistent with metabolic scaling alone, as predicted by the Yates et al. (2025) f(T) framework; a field slope exceeding physiological limits would indicate that ecological processes inflate it. The study builds directly on Ogonowski (2026), in which field catch patterns indicated that night duration, rather than temperature, was the principal predictor of light-trap catches of stickleback in these same bays; this supports their use as a largely temperature-independent abundance reference. It extends the Yates et al. (2025) metabolic framework by delivering the first field-based characterisation of f(T) for eDNA production across a continuous seasonal gradient, anchored by independent abundance benchmarks from two gear types with opposing temperature dependencies.

## 2. Methods

### 2.1. Study site and sampling design

Sampling was conducted in four shallow, semi-enclosed coastal bays at Askö island in the southern Stockholm archipelago, Sweden (58°49’N, 17°39’E), during the spring and early summer of 2023. The bays are characterised by salinities of approximately 7 psu, shallow depths (0.5–2 m), and extensive littoral habitat suitable for *G. aculeatus* spawning and foraging. Water temperature was recorded weekly in each bay using a Rinko ASTD-102 profiler (JFE Advantech Co., Ltd, Japan). The median value was calculated per depth profile and across transects to provide a grand median per bay visit (range: 5.7–20.5°C across the season).

Each bay was treated as an independent replicate. On each of eight sampling occasions (late April to early July), we deployed both light traps and benthic fish traps within each bay and collected eDNA water samples on the same day as trap retrieval or within two days thereafter, consistent with eDNA persistence times in coastal brackish water (Collins et al. 2018).

Quatrefoil light traps were deployed overnight at fixed stations within each bay (*n* = 5 per bay) equipped with green LED lights to attract fish by phototaxis. The number of adult *G. aculeatus* per trap was recorded at retrieval. Collapsible trap nets (45 × 24 × 24 cm, 3 mm mesh; Ogonowski, 2026) were deployed simultaneously with light traps at the same fixed stations. These traps operate without attractants and rely on fish simply encountering and entering the trap.

Two eDNA transects (one following the vegetated coastline and the other in the deeper central part of the bay) were sampled per bay and sampling date. In total, 88 eDNA samples were collected across the season: 64 were matched to concurrent trap deployments and entered the main analysis, while 16 pre-season samples and 8 post-season samples collected after the final trap deployment were retained as unmatched observations to document the full seasonal trajectory of the eDNA signal outside the matched sampling window (Figure S 3). The study design and sampling infrastructure are described in detail in Ogonowski (2026).

### 2.2. eDNA extraction and quantification

Water samples were collected along two fixed transects within each bay following the protocol described in Ogonowski et al. (2023). One litre of surface water was collected at 50 m intervals along each transect and filtered immediately in the field using enclosed eDNA Dual Filter Capsules (0.8 µm polyethersulfone membrane; Sylphium Molecular Ecology, Groningen, the Netherlands; SYL009) with a peristaltic pump and sterile silicone tubing.

Following filtration, a preservation/lysis buffer (Sylphium Molecular Ecology, Groningen, the Netherlands) was injected directly into the capsule, which was then sealed with Luer lock caps according to the manufacturer’s protocol. Field negative controls consisted of one litre of DNA-free MilliQ water filtered per sampling event. Prior to sampling, the boat was decontaminated with a 10% household bleach solution to minimise cross-contamination risk from prior fish handling. Environmental DNA was subsequently isolated using the Sylphium eDNA Isolation Kit (Sylphium Molecular Ecology, Groningen, the Netherlands; SYL002) according to the manufacturer’s protocol. Disposable nitrile gloves were used at all times while handling eDNA equipment and samples.

We used a real-time, species-specific quantitative polymerase chain reaction assay (qPCR) to quantify three-spined stickleback DNA in collected samples. The primer and probe combination (F-primer: 5′-ACGCCACCTTAACACGTTTC-3′, R-primer: 5′-AGAGCCTGTCTGGTGAAGGA-3′, and probe: 5′-FAM-CTGGTGCCACACTTGTTCAC-MGB-3’; Microsynth AG) was originally developed by Thomsen et al. (2012) and targets a 101 base-pair fragment of the cytochrome b gene (cytb). qPCR was performed on a BioRad CFX384 Real-time PCR system with 15 µL reaction volumes in quadruplicate (technical replicates) for each sample on the FAM channel. To control for potential inhibition at the level of the DNA extract, we used an internal positive control (IPC, Cy®5-QXL®670 Probe; EuroGentec) co-amplified in duplex with two northern pike (*Esox lucius)* assays (Elu_COI and Elu_5s_rDNA) on the Cy5 channel. These reactions were run on the same extracted DNA as the *G. aculeatus* assay, but in separate wells. The *G. aculeatus* assay was run as singleplex on the FAM channel. The IPC therefore detects inhibition in the DNA extract but not inhibition specific to the *G. aculeatus* reaction. eDNA was quantified using a standard curve consisting of an 8-step, 10-fold dilution series of stickleback DNA extracted from muscle tissue with a Chelex protocol (Karlsson et al. 2022). 4 µL of template DNA was used in each reaction. Thermal cycling consisted of an initial denaturation at 95°C for 10 min, followed by 50 cycles of 95°C for 15 s and 60°C for 60 s.

The standard curve was estimated by maximum likelihood, jointly fitting amplification efficiency, residual Cq standard deviation, and the absolute DNA-to-molecule scaling constant, with non-detects accommodated via a Poisson-mixture likelihood (Schmidt et al. 2023). Per-sample measurement error was propagated to downstream models as a known response standard error.

Non-detects at the lowest standard dilution step (6 of 8 replicates failed to amplify) reflect expected Poisson sampling variation when DNA molecules are rare. At these extreme dilutions the log-linear standard curve predicts Cq values beyond the instrument’s observable range (50 cycles), so extrapolating it to obtain a limit of detection (LOD) is not valid. Instead, the LOD was estimated by anchoring to the empirical detection rate at the lowest dilution step and scaling to the 95% detection probability criterion using the Poisson-mixture model, with a correction for the selection bias that arises because only detected replicates contribute to the mean Cq at that step (Forootan et al. 2017, Lesperance et al. 2021). This yielded LOD = Cq 38.80. Quantification precision across the working range, assessed as the coefficient of variation on the log-concentration scale (logCV = 15.2; Forootan et al. 2017, Klymus et al. 2020), satisfied the 35% criterion throughout, so the limit of quantification (LOQ) equals the LOD.

The eDNA response variable was defined as edna_index = −mean_Cq, so that the index increases with eDNA concentration; this sign convention ensures all model coefficients are interpretable in the positive direction. Sampling effort corrections for filtered volume and bay area (dilution) were applied as log-transformed offset terms. Three negative control types (field blanks, extraction blanks, PCR blanks), inhibition assessment via the IPC, and the evaluation of flagged samples are reported in the Supplementary QC section S2.

### 2.3. Data handling and statistical analyses

We analysed the data in four steps (Figure 1), of which two are the primary modelling stages referred to throughout as Stage 1 and Stage 2. First, as a diagnostic, raw correlations among eDNA, light-trap, and benthic-trap catches established whether the opposing temperature sensitivities of the two gears produced the predicted signal structure before any correction. Second, gear-specific Bayesian GLMMs standardised trap catches for their primary observation-process biases, yielding the combined abundance index. Stage 1 then characterised the eDNA–temperature relationship across six candidate functional forms; residuals from the best-supported model served as the empirically corrected eDNA predictor, while a parallel model with the temperature slope constrained to the laboratory maximum metabolic rate (MMR) yielded the first-principles (FP) corrected predictor, applied identically to the benthic trap. In Stage 2, the combined abundance index was modelled as a function of each of three eDNA predictors — raw, FP-corrected, and empirically corrected — in three otherwise identical Bayesian linear models, enabling direct comparison of the eDNA–abundance slope across correction strategies. All Bayesian models were fitted using brms (Bürkner 2017) with Stan (Carpenter et al. 2017). Prior sensitivity results and full model specifications are provided in Supporting Information. All analyses were conducted in R 4.5 (R Core Team 2025).

**Figure 1.**
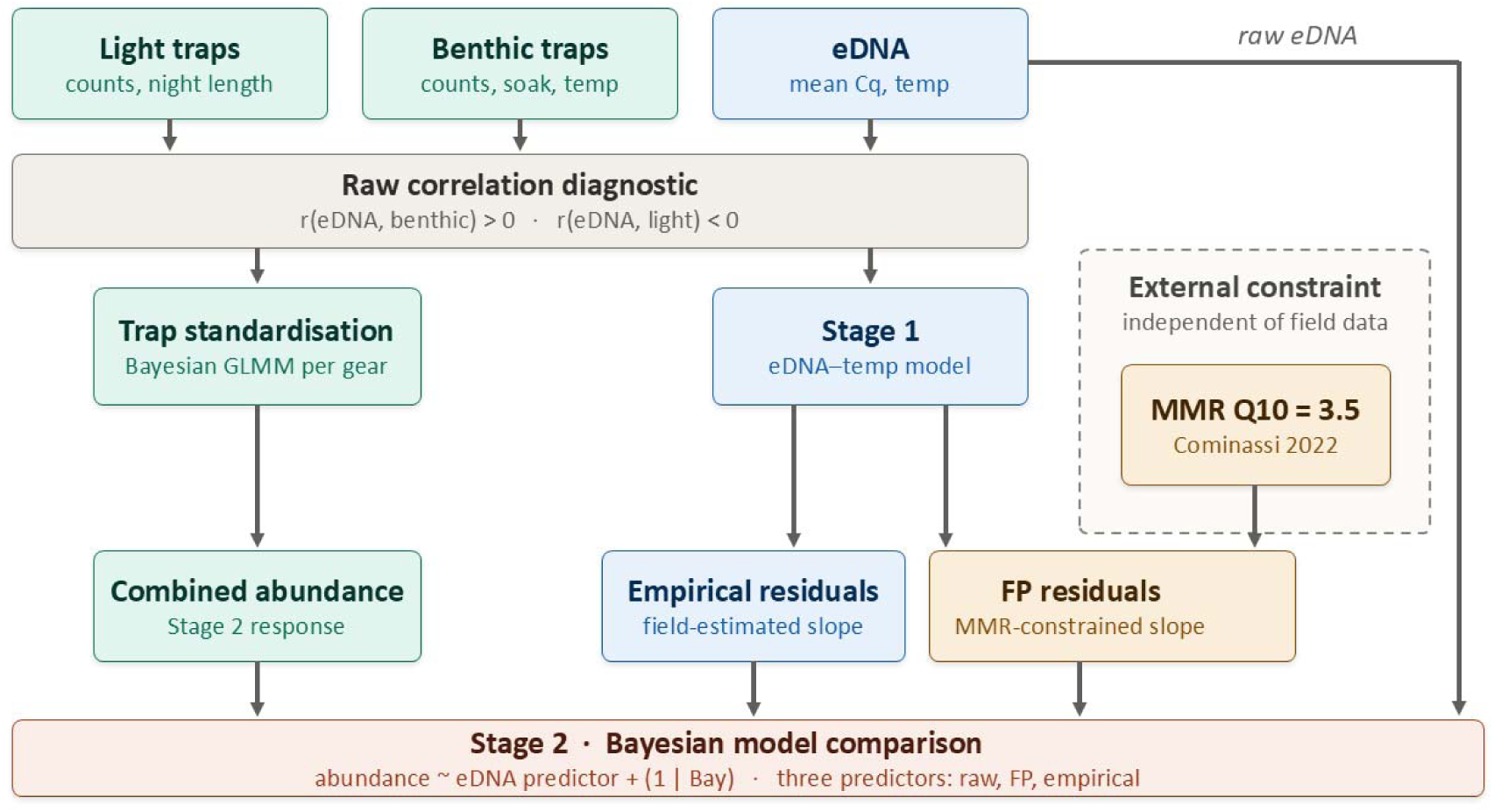
Schematic flow-chart of the statistical analysis pipeline.

#### 2.3.1. Trap catch standardization

Light-trap catches were standardised using a Bayesian generalised linear mixed model with centred night duration as a predictor, because light-trap catchability depends on the length of the night over which the light source attracts fish. Model-predicted catches at mean night duration constituted the night-corrected index *lt_z,* expressed as within-bay z-scores. To test whether abundance changed systematically across the season, *lt_z* was regressed on day-of-year in a Bayesian linear mixed model with bay as a random intercept, using the light-trap index as the most temperature-independent abundance proxy available.

Benthic-trap catchability depends strongly on fish activity, which is governed by temperature (Sullivan 1954, Rudstam et al. 1984). Rather than estimating this temperature dependence empirically from the field data, which would conflate metabolic activity with the seasonal abundance change we sought to recover, we applied a first-principles physiological constraint: soak time was scaled into an effective metabolic soak time using the species-specific maximum metabolic rate Q10 (3.5; Cominassi et al. 2022), and this effective effort entered the model as an offset, with no free temperature predictor. Model-predicted catches at the reference temperature constituted the temperature-corrected index *bt_z*, expressed as within-bay z-scores.

Because neither method was optimal throughout the survey period, with light traps becoming ineffective during the midsummer period when high-latitude photoperiod leaves no true darkness, we combined the two indices using a precision-weighted average in which each gear contributed in inverse proportion to its estimation variance at each bay and date. This downweighs a gear when its index is poorly constrained, for example light traps as catches approach zero in midsummer, allowing the metabolism-corrected benthic index to dominate the high-summer trajectory without arbitrary truncation. Full model specifications, prior justifications, and the weighting framework are given in Supporting Information S1.1.

#### 2.3.2. Stage 1 — eDNA–temperature model and first-principles correction

As an initial test of whether temperature dominates the eDNA signal, Pearson correlations were computed between raw eDNA concentration and raw catches from each trap type, and between the two trap types. Six candidate models were then fitted to the eDNA data, combining three functional forms (linear, log-linear, quadratic) with and without a bay-area offset, and compared via Pareto Smoothed Importance Sampling Leave-One-Out (PSIS-LOO) cross-validation; models within 1 SE of the top model were considered equivalent (Vehtari et al. 2017). Among statistically equivalent forms, the log-linear form was preferred on mechanistic grounds, because log-linear temperature scaling maps directly onto Q10/Arrhenius metabolic rate theory and allows the temperature coefficient to be interpreted as a physiological scaling exponent. From the selected model, the temperature coefficient was back-transformed to a natural-scale slope (*b_raw*) and converted to a field Q10 evaluated at the mean observed temperature (*T_ref* = 12.5°C), following the Q10 definition of Yates et al. (2025). Because the model uses log-linear temperature scaling, Q10 is range-dependent, and the reference temperature must be specified for comparability with laboratory benchmarks. Posterior mean residuals from the empirical model served as the empirically corrected eDNA predictor in Stage 2.

A parallel model was fitted with the temperature slope constrained to MMR Q10 = 3.5 via a tightly informative prior (SD = 0.001), yielding the FP-corrected eDNA predictor. MMR represents the physiological ceiling on sustained temperature-driven metabolic activity in ectotherms (Clarke and Johnston 1999, Norin and Clark 2016); any field Q10 exceeding this value therefore reflects ecological processes — abundance change or behavioural amplification beyond metabolism alone. Constraining the correction to the MMR ceiling ensures that only the metabolic component of the temperature signal is removed, leaving ecological variance intact. The MMR Q10 of 3.5 corresponds to the upper bound of the range reported for *G. aculeatus* at 5–12°C (Cominassi et al. 2022) and was chosen as the most physiologically defensible ceiling for this species and temperature range.

#### 2.3.3. Bayesian model comparison: eDNA as abundance predictor

Prior to model fitting, eDNA predictors were aggregated to bay-date means (n = 32; 4 bays × 8 dates), the lowest level of observation shared by all variables, because the abundance index is measured at the bay-date level. The within-bay-date intraclass correlation coefficient (ICC = 0.678) confirmed that the bay-date mean captures the dominant component of transect-level eDNA variation.

We then fitted three Bayesian linear models in which the GLMM-corrected combined trap abundance index was modelled as a function of a standardised eDNA predictor, with bay as a random intercept to account for repeated measures across eight sampling dates within each of the four bays:

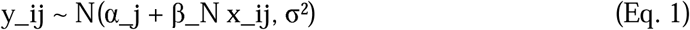

where y_ij is the combined abundance index for date i in bay j; x_ij is the standardised eDNA predictor; β_N is the eDNA–abundance slope; and α_j is the bay-level random intercept. The three models differ only in the definition of x_ij: volume-corrected raw eDNA (no temperature correction), FP-corrected eDNA residuals (temperature slope constrained to MMR Q10), and empirically corrected eDNA residuals (temperature slope estimated from field data). All models used weakly informative priors (β_N ∼ N(0,1); intercept ∼ N(0,2); σ ∼ Exponential(1); σ_α ∼ Exponential(1)) and were fitted with 4 chains × 4000 iterations (1000 warmup) at adapt_delta = 0.99.

## 3. Results

### 3.1. Raw three-way correlation structure between traps and eDNA

The raw seasonal trajectories (Figure 2) reveal opposing dynamics across all four bays: raw eDNA and benthic-trap catches both rise across the season in parallel with temperature, while light-trap catches decline as nights shorten. The raw correlations confirmed this opposing-signal pattern — eDNA correlated positively with raw benthic-trap catches (r = 0.434, p = 0.01) and negatively with raw light-trap catches (r = −0.599, p < 0.001), while the correlation between the two trap types was not statistically significant (r = −0.216, p = 0.235). This structure is consistent with eDNA and benthic-trap catches sharing the same underlying thermal-activity mechanism, while light-trap catchability operates on an opposing seasonal axis governed by night duration. After standardising each gear type for its primary bias, both abundance indices follow broadly consistent trends across bays (Figure 3), confirming that the opposing raw patterns reflect observation-process confounds rather than true abundance differences.

**Figure 2.**
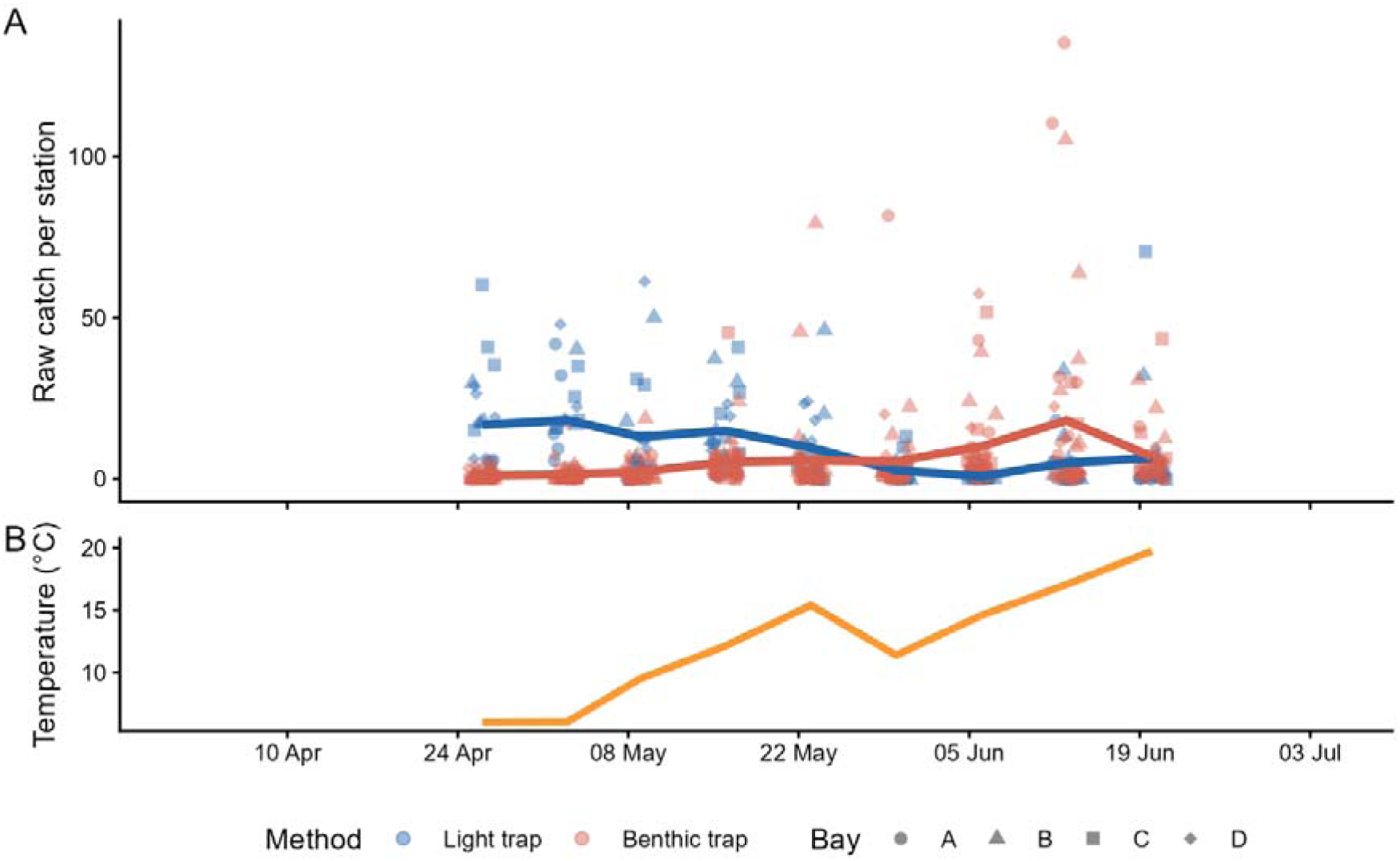
Raw seasonal trajectories of light-trap (blue points and mean line) and benthic-trap (red points and mean line) station catches (numbers per trap) to show variation per method and sampling date (A), and corresponding mean water surface temperature (B, orange line).

**Figure 3.**
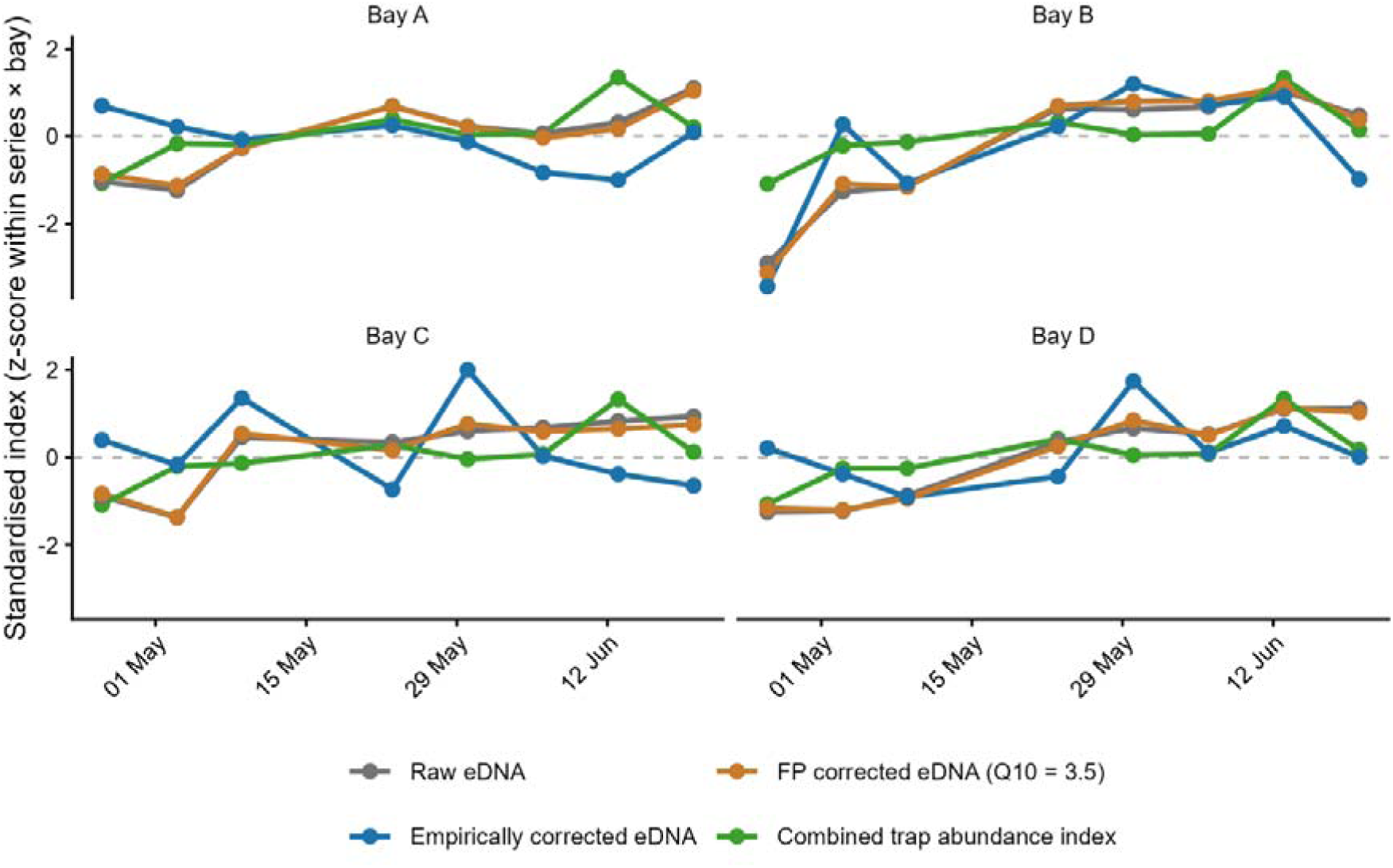
Standardised seasonal trajectories of eDNA and the combined trap abundance index in the four study bays (A–D), shown as within-bay z-scores against sampling date. Series are raw eDNA (grey), eDNA after empirical temperature correction (blue), eDNA after first-principles temperature correction constrained to the maximum metabolic rate Q10 of 3.5 (orange), and the combined trap abundance index (green). Each eDNA point is the bay-date mean of the two transect samples. Points show one value per bay and sampling date, connected by straight segments between consecutive dates. The dashed horizontal line marks the within-bay mean (z = 0).

### 3.2. Standardized abundance indices from traps

The night-corrected light-trap index showed a positive but statistically unresolved temporal trend (β = 0.010 per day, 95% CrI: −0.007 – 0.026; posterior probability of a positive slope 0.88). Because light-trap catchability is predominantey driven by night duration rather than temperature (Ogonowski 2026), this index is the most temperature-independent abundance indicator available in this system. Its positive point estimate is consistent with a seasonal increase in stickleback abundance, but the trend is not statistically resolved, so we cannot separate a genuine abundance increase from thermal and behavioural amplification of shedding on the basis of this signal alone.

The combined abundance index, based on standardised catches from both light and benthic traps across the full season, showed a clearer positive temporal trend (β = 0.025 per day, 95% CrI: 0.016 – 0.033). This is consistent with an increase in abundance over the sampling season

(Figure 3).

### 3.3. Stage 1: temperature dominates the eDNA signal

Leave-one-out-cross-validation (LOO-CV) comparison ranked the quadratic and log-linear forms (without area offset) as statistically equivalent (ELPD difference = 1.5 ± 1.9 SE); the log-linear form was retained for the reasons given in Methods 2.3.2. Temperature explained 65% of the variance in eDNA (Bayesian R² = 0.654, 95% CrI: 0.544–0.715), and the back-transformed thermal scaling exponent was *b_raw* = 4.28 (95% CrI: 3.48–5.07). Full LOO-CV results are provided in Table S 1.

### 3.4. Thermal scaling diagnostics: field Q10 far exceeds metabolic limits

The field Q10 for eDNA substantially exceeded laboratory metabolic benchmarks (Figure 4). Evaluated at mean temperature (12.5°C), the eDNA Q10 was 12.4 (full season), well above the MMR Q10 range of 1.5–3.5 for *G. aculeatus* (Ressel et al. 2022, Cominassi et al. 2022). Prior sensitivity results are reported in the Supporting Information section S1.3.

**Figure 4.**
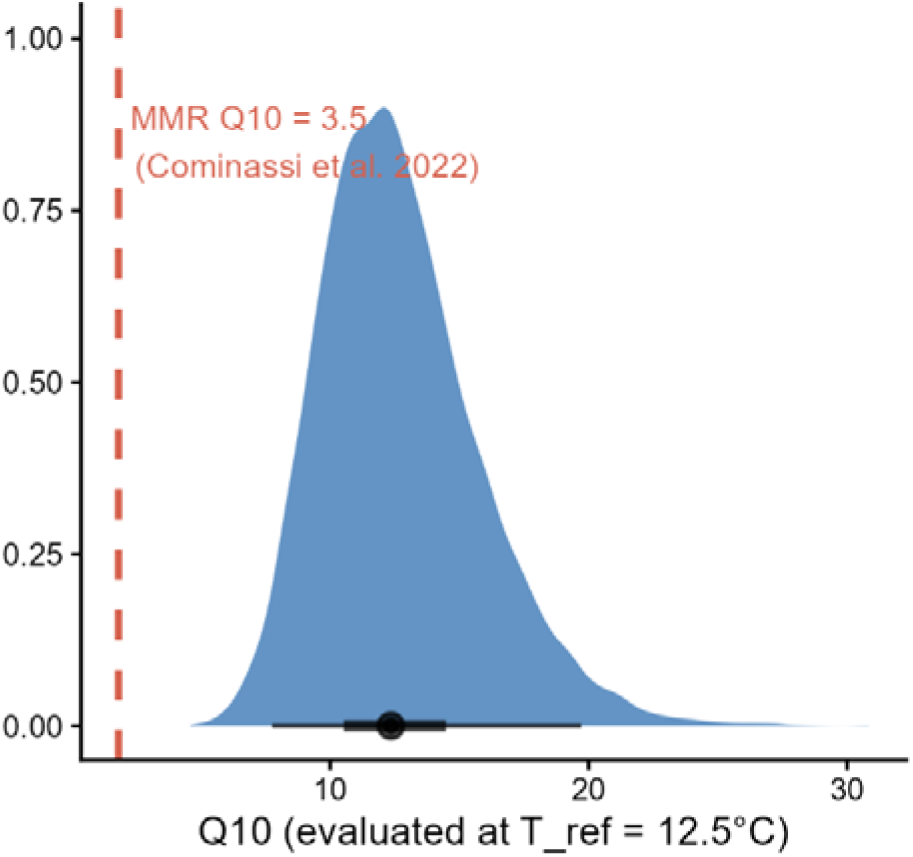
Posterior distribution of the field-derived eDNA Q10 coefficient evaluated at a reference temperature of 12.5°C (blue density plot). Point and whiskers show the median and 50–95% CrI. The dashed red line indicates the MMR Q10 calculated from metabolic rate measurements at 5–12°C (Q10 = 3.5; Cominassi et al., 2022).

### 3.5. Bayesian model comparison: eDNA as an abundance predictor

The three Bayesian models showed a clear ordering (Figure 5). Raw eDNA (no temperature correction) predicted the GLMM-corrected abundance index strongly (β_N = 0.451, 95% CrI: 0.286 – 0.617). FP-corrected eDNA (temperature slope constrained to MMR Q10 = 3.5) performed almost identically (β_N = 0.430, 95% CrI: 0.259 – 0.605). Empirically corrected eDNA (temperature slope estimated from field data), by contrast, showed no credible abundance relationship (β_N = 0.090 95% CrI: −0.157 – 0.328), with 24% of the posterior lying below zero. This ∼5-fold difference in β_N between empirically corrected eDNA and both raw and FP-corrected eDNA shows that empirical temperature correction in a seasonal system suppresses the abundance signal to noise level, whereas raw and FP-corrected eDNA perform almost identically.

**Figure 5.**
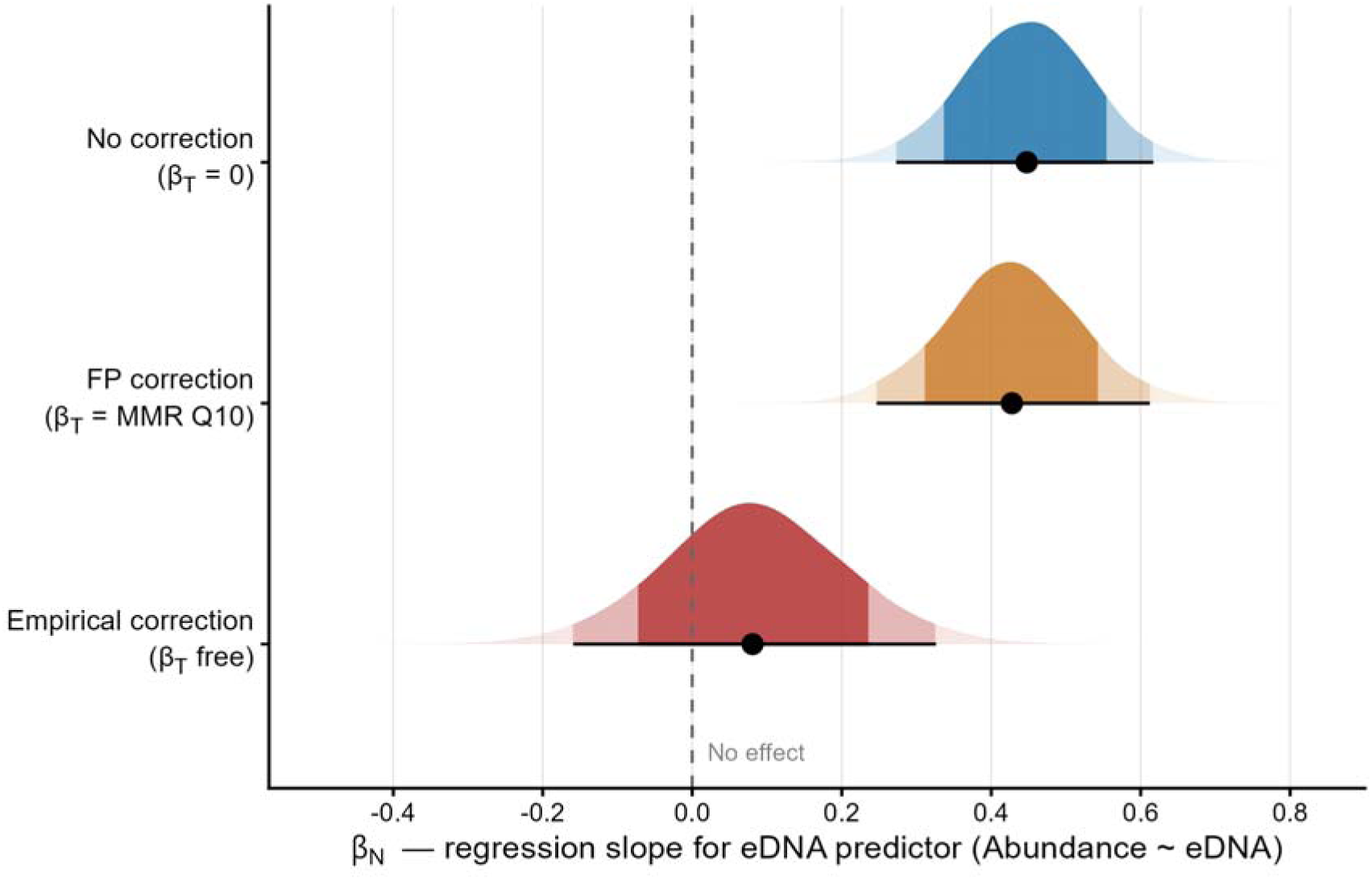
Posterior distributions of the regression slope β_N (combined abundance index ∼ standardized eDNA) from three Bayesian linear models differing only in how the eDNA predictor is temperature-corrected: no correction (β_T = 0; blue), FP correction (β_T = MMR Q10; amber), and empirical correction (β_T estimated from data; red). Inner and outer bands show 80% and 95% posterior credible intervals; the point indicates the posterior median.

## 4. Discussion

In temperate ecosystems, seasonal eDNA monitoring faces a fundamental identification problem when temperature drives both eDNA dynamics and the catchability of conventional passive gears. Consequently, any apparent eDNA–abundance relationship may reflect a shared thermal response rather than a true density signal. We addressed this unidentifiability problem by deploying two trap types alongside eDNA: a light trap driven by photoperiod, which provides a largely temperature-independent abundance reference, and a benthic trap whose temperature dependence shares the same locomotor-metabolic basis as the production term governing eDNA concentration. This triangulation isolates the thermal component of the eDNA signal and tests whether it tracks abundance once that component is physiologically constrained. Triangulating these signals lets us quantify the thermal component of the eDNA signal against an independent physiological benchmark and test whether eDNA tracks abundance once that component is constrained rather than empirically removed.

### 4.1. The field temperature signal exceeds metabolic predictions

Estimating the temperature effect on eDNA from field data yielded a Q10 of 12.4, well above the MMR Q10 range of 1.5–3.5 reported for three-spined stickleback across laboratory temperature treatments (Ressel et al. 2022, Cominassi et al. 2022), confirming that the seasonal eDNA slope reflects a compound of metabolic shedding, behavioural amplification, and abundance change that cannot be attributed to temperature alone. A substantial part of the behavioural component is likely reproductive: elevated eDNA shedding during spawning, from gamete release, nest-building, agonistic interactions, and spawning aggregations, is well documented (Takeuchi et al. 2019, Ostberg and Chase 2022), and our sampling period spans the Baltic stickleback breeding season (Borg 1985). That the raw eDNA signal nonetheless retains a detectable abundance association reflects the positive covariance between temperature and abundance in this system, not a general property of uncorrected eDNA.

This excess has a direct implication for the temperature dependency function f(T), the metabolic production term formalised by Yates et al. (2021) and extended by Yates et al. (2025). Yates et al. (2025) proposed that f(T) might be characterised from field data spanning a sufficient temperature gradient; our results show why this is not achievable in a seasonal system. Because eDNA production cannot be separated from abundance and behaviour when all three covary with temperature, a field-estimated slope quantifies the total temperature dependence of the signal, not f(T) itself. This distinction has practical consequences: in a simulation that explicitly modelled temperature-dependent eDNA production, Jo et al. (2025) found that the sensitivity of the eDNA–abundance relationship was preserved only when the production mechanism was modelled, and degraded when it was ignored. Our field data provide the empirical counterpart, demonstrating that the slope recoverable from seasonal field sampling conflates production with ecological change and therefore cannot be used as f(T).

### 4.2. First-principles correction preserves the abundance signal

The Bayesian model comparison showed not only that FP correction outperformed empirical correction, which was expected, but that raw eDNA performed as well as FP-corrected eDNA. This equivalence arises because temperature and abundance increased together across the season. This co-variation makes the temperature-driven inflation of eDNA act in the same direction as the abundance signal, so raw eDNA overestimates absolute concentration at high temperature while its association with abundance, measured by the abundance-regression slope β_N, remains strong. Because the MMR-constrained slope is shallow, FP correction removes only a small part of the eDNA variation and leaves this association largely intact.

The equivalence between raw and FP-corrected eDNA is specific to systems where temperature and abundance covary positively, as they did here. Where this covariance does not hold, raw eDNA would be confounded. This includes systems where abundance peaks in cold water, where populations are stable while temperature varies, or where reproductive shedding decouples eDNA from density. A stable pearl mussel population, for instance, produced a roughly 20-fold seasonal swing in eDNA driven by factors other than abundance (Wacker et al. 2019).

In such systems, a physiologically constrained correction can recover abundance that raw eDNA does not. Beaulieu et al. (2026) showed this by combining bioenergetics and mass-balance modelling, raising the variance explained in eDNA from 24% to 71% against mark-recapture estimates. The value of FP correction here is therefore not that it improved on raw eDNA but rather that it remains reliable under temperature–abundance relationships that in most cases cannot be known beforehand.

Empirical correction, by contrast, causes biased abundance estimates precisely when abundance changes seasonally alongside temperature. Under such conditions, the field slope absorbs the abundance signal together with the metabolic component of eDNA production.

### 4.3. Water residence time and the strength of temperature confounding

The exceptionally steep temperature dependence observed in our field data (Q10 = 12.4) is partly a structural property of our open coastal study system, driven by water residence time.

To understand how hydrodynamic environments modulate the eDNA temperature signal, two distinct physical mechanisms must be uncoupled: signal magnitude and signal sensitivity.

#### 4.3.1. Signal magnitude vs. thermal sensitivity

The first mechanism concerns absolute signal magnitude. High water exchange rates and large basin volumes dilute local eDNA concentrations. While this dilution dictates the baseline concentration of detectable DNA, it does not inherently alter the slope of the temperature response. This explains why our Stage 1 models including a static bay-area dilution offset did not outperform simpler variants. In dynamic coastal environments, static volume is a poor proxy for dilution; concentration gradients are instead governed by the dynamic balance between local degradation kinetics and hydrodynamic flushing.

The second, more critical mechanism acts directly on the apparent temperature sensitivity (Q10) of the eDNA signal. In a closed water body at a structural steady-state, eDNA concentration is a function of the ratio between the shedding rate (r_S_) and the degradation rate (r_D_):

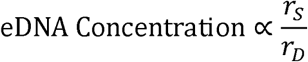

Because both cellular shedding and microbial degradation accelerate with warming, their respective temperature dependencies partially cancel each other out in the numerator and denominator. Consequently, the net temperature sensitivity of measured eDNA in closed systems is muted relative to the true temperature dependence of shedding alone.

#### 4.3.2. Open systems inflate the temperature effect

In an open coastal system characterized by rapid water exchange, this functional balance shifts fundamentally. Here, the primary loss term shifts from temperature-dependent biological degradation (r_D_) to temperature-independent hydrodynamic flushing (r_F_), driven by wind, waves, and coastal currents:

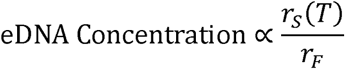

This hydrodynamic mechanism explains why the metabolic and behavioral components of our stickleback eDNA signal were so heavily amplified. Recent lagrangian particle transport simulations and tidal field studies support this pivotal role of hydrodynamics, confirming that water residence times dictate whether decay kinetics or physical transport dominate the local eDNA-abundance relationship (Clark et al. 2025, Zanni et al. 2025). In flushing-dominated environments like our coastal bays, hydrodynamic transport strips away the thermal buffering provided by degradation, exposing a highly sensitive, temperature-confounded signal that biases abundance inferences if left uncorrected.

### 4.4. Abundance is not stable: resolving the identifiability question

A central question in any seasonal eDNA study is whether rising eDNA concentrations reflect genuine abundance change or merely thermal amplification of shedding from a stable population. Our two trap-based abundance indices address this question but cannot fully resolve it. The night-corrected light-trap index, the most temperature-independent abundance proxy available here, showed a positive but statistically unresolved seasonal trend. On its own, this index could not unequivocally demonstrate that abundance increased, in part because light-trap catches fell to near zero around midsummer, when the absence of true darkness at this latitude leaves the night-duration correction unable to recover a usable signal. The light traps therefore effectively stopped tracking abundance during a period when stickleback densities would be expected to be high –in the Baltic, stickleback breeding extends from late April to late June or early July (Borg 1985), when adults migrate from the open sea to aggregate at high density in shallow coastal bays (Bergström et al. 2015).

The combined index, which draws on both light and benthic traps and so bridges the midsummer period when the light traps fail, showed a clear positive seasonal trend. This index is not fully independent of temperature, because its benthic component is corrected for rather than free of thermal effects, so it provides a less stringent test of temperature-independence than the light traps alone. Nonetheless, its agreement with the positive point estimate of the light traps raises the probability that abundance genuinely increased over the season. We therefore interpret the seasonal eDNA rise as reflecting a real increase in stickleback abundance in addition to thermal and behavioural amplification of shedding, while acknowledging that these two contributions cannot be fully separated with the present data.

### 4.5. Implications for monitoring design

Three practical recommendations emerge from this study. First, before interpreting any eDNA–abundance correlation, practitioners must evaluate whether the reference method shares observation-process biases with eDNA. In our system, the uncorrected light trap produced a strongly negative correlation with eDNA (r = -0.599) because the reference was confounded by night duration. A practitioner relying on this single correlation would wrongly conclude that eDNA is a poor abundance indicator. This failure mode is a broad risk: gillnet and electrofishing catchability are strongly temperature-dependent (Olin et al. 2016, Simonson et al. 2022). Where temperature correction of eDNA is applied, physiologically constrained corrections using species-specific maximum metabolic rate MMR Q10 values should be preferred over empirical field-derived slopes.

Second, deploying two trap types with opposing environmental drivers provides the most robust validation framework. The temperature-independent method establishes the abundance trend, the temperature-dependent method confirms the shared observation process, and together they yield a combined index suitable for physiologically grounded eDNA calibration. The performance of our combined index (β_N = 0.448 for raw eDNA, Pearson r = 0.70) is comparable to published data in coastal and riverine systems (Nakagawa et al. 2022, Morrison et al. 2023), confirming that appropriately corrected multi-method designs can recover meaningful abundance signals even in open, hydrodynamically complex environments.

Third, the first principles-framework applies broadly to any species for which laboratory respirometry data exist. The growing bioenergetics literature makes this feasible for many commercially and ecologically important fish species, and the Yates et al. (2025) framework provides the structural scaffolding to integrate these metabolic parameters into standardised monitoring protocols.

## 5. Conclusion

In seasonal aquatic systems, temperature variations dominate raw environmental DNA signals. This study demonstrates that the methodology used to account for thermal dependence determines whether an absolute abundance signal can be recovered or not. While empirical field-derived corrections remove the true density signal due to seasonal temperature-abundance covariance, a correction constrained to laboratory-derived metabolic rates successfully preserves it.

We recommend that physiologically constrained corrections serve as the default for seasonal eDNA monitoring programs. The FP framework matches the tracking efficiency of raw eDNA under positive environmental covariance while safeguarding against signal loss when density and temperature decouple. Ultimately, validating quantitative eDNA frameworks requires independent reference gears whose underlying observation-process biases are decoupled from the environmental sensitivities of eDNA shedding. These two foundational principles provide a rigorous, reproducible path toward quantitative eDNA-based abundance assessments across temperate coastal marine ecosystems.

## 6. Acknowledgements

We would like to thank Johan Wenngren (Askö, Sweden) and associates for carrying out the fish sampling and technical assistance; AquaBiota Water Research for conducting the eDNA sampling; and the Stockholm University Baltic Sea Centre, Askö Laboratory for providing housing and equipment. We also thank Erik Karlsson (Swedish University of Agricultural Sciences) for technical assistance with the laboratory DNA extractions and PCR preparation, and Björn Rogell (Swedish University of Agricultural Sciences) for valuable comments on the manuscript.

## 8. Author contributions

Conceptualization: M.O; Methodology: M.O; Data curation: M.O; Formal analysis: M.O, Z.G; Funding acquisition: M.O, Project administration: M.O; Visualization: M.O; Writing – original draft: M.O; Writing – review & editing: M.O, Z.G.

## 9. Ethical statement

All applicable international, national, and/or institutional guidelines for the care and use of animals were followed. The fish sampled and handled in this study complied with the standards and procedures stipulated by the Swedish Ministry of Agriculture and the Swedish Board of Agriculture. The research was conducted under a formal permit for animal experimentation, which was approved by the Linköping Regional Animal Ethics Committee (DNR: 23379-2022). All measures possible were taken to follow the 3R tenets (Replacement, Reduction, and Refinement) as part of the ethical vetting process for our permit application. Specifically, refinement was achieved through the use of passive, non-lethal gear (light traps and fish traps). The study followed the taxon-specific guidelines for the ethical treatment of fishes as outlined by the Stockholm University Baltic Sea Centre (document number: SU-484-0009-22). Following identification and quantification, all bycatch was monitored for normal swimming behaviour and released immediately at the site of capture.

## 10. Data sharing

All relevant data and R-scripts used in this manuscript are available through Figshare: https://doi.org/10.6084/m9.figshare.32834141.

