## Supporting Information for "Field-derived temperature correction compromises eDNA-based abundance inference"

Supplementary Information for: Field-derived temperature correction compromises eDNA-based abundance inference

S1 Supplementary methods and results

This section provides full specifications for the statistical models used in the main analysis, together with convergence diagnostics and prior justifications.

S1.1 Trap catch standardisation models and prior justification
     S1.1.1 Light trap standardization

Light-trap catches were modelled with a negative binomial generalised linear mixed model (GLMM, log link), appropriate for overdispersed count data. Mean centred night duration (hours) and bay entered as fixed effects, while date and station entered as crossed random intercepts to absorb among-occasion and among-station heterogeneity. Because night duration was centred, the intercept corresponds to the expected log-count at mean night duration and the slope to the change in log-count per additional hour of darkness, allowing weakly informative priors to be set on an interpretable scale (Gelman et al. 2008, 2017). The intercept received a Normal(0, 5) prior, reflecting broad uncertainty over plausible baseline catch magnitudes on the log scale. Population-level coefficients were assigned Normal(0, 2) priors, permitting a wide range of effect sizes while regularising against implausibly large estimates. Random-effect standard deviations were given Exponential(1) priors, concentrating variance components toward zero following standard conventions. The negative binomial shape parameter received a Gamma(2, 0.5) prior, permitting substantial overdispersion while avoiding values near zero.

S1.1.2 Benthic trap first-principles standardization

To prevent the temporal collinearity between spring water warming and spawning migration from biasing the density signal, benthic-trap catches were standardized using a first-principles ecophysiological constraint rather than an empirical temperature covariate. Soak time (hours) was scaled into effective metabolic soak time to account for temperature-dependent locomotor activity and gear encounter probability (Rudstam et al. 1984, Killen et al. 2015). Using a species-specific active metabolic ceiling derived from laboratory respirometry ($Q_{10}=3.5$, evaluated at a baseline reference temperature $T_{\text{ref}}={12.5}^{\circ}\text{C}$; Cominassi et al., 2022), effective effort was computed as:

$$\text{Effort}_{\text{effective}}=\text{Soak Time (h)}\times{3.5}^{\frac{T_{\text{field}}-12.5}{10}}$$

The natural logarithm of this effective metabolic effort entered the model as a fixed offset term. No free temperature predictor was estimated. The model structure (fixed bay intercepts and crossed random intercepts for date and station) and the prior framework remained identical to the light-trap formulation. The benthic response variable was the station-level catch per bay-date, rounded to the nearest integer for the negative binomial likelihood.

S1.1.3 Construction of the combined abundance index

Because the nights are very short during midsummer (June–July) at high-latitudes, the light traps stop working when the structural absence of dark nights reduces catchability ($q_{L}\to0$). To seamlessly transition between the complementary gears, station-level posterior draws for both metrics were extracted on the log-linear predictor scale using the posterior *linpred* function. Draws were aggregated to bay-date matrices to calculate the joint posterior mean and estimation variance for each sampling occasion.

The two independent standardized indices (*lt_z* and *bt_z*) were then combined into a single unified density baseline (the combined abundance index) using an inverse-variance precision weighting framework at each bay-date timestep (*t*):

$$\text{Index}_{\text{joint},t}=\frac{\frac{lt_{z,t}}{\sigma_{lt,t}^{2}}+\frac{bt_{z,t}}{\sigma_{bt,t}^{2}}}{\frac{1}{\sigma_{lt,t}^{2}}+\frac{1}{\sigma_{bt,t}^{2}}}$$

By conducting this weighting on the log-linear scale, the high uncertainty generated as light-trap catches approach zero automatically deflates the gear's weight ($\frac{1}{\sigma_{lt}^{2}}\to0$), allowing the metabolism-corrected benthic baseline to dominate the high-summer index without requiring arbitrary manual truncation thresholds.

S1.1.4 Model implementation and diagnostics

All Markov Chain Monte Carlo (MCMC) simulations were implemented in Stan via the brms package (Carpenter et al. 2017, Bürkner 2017) using 4 parallel chains running for 4,000 iterations each, with a 2,000-iteration warmup phase. Convergence and sampling efficiency were verified using the Gelman-Rubin diagnostic ($\hat{R}\leq1.01$across all parameters), visual inspection of chain trace plots for proper mixing, and ensuring sufficient effective sample sizes ($ESS>1,000$).

S1.2 Stage 1 model specification and prior justification

The Stage 1 model estimated the eDNA–temperature relationship across six candidates, combining three functional forms for f(T) (linear, log-linear, and quadratic) with and without a bay-area offset to account for dilution of eDNA. Each was fitted as a Gaussian mixed model with a measurement-error response (Eq. S1–S3). The eDNA index for observation *i* in bay *j*, y_{ij}, was modelled with a total variance equal to the sum of an estimated residual variance, σ², and the known per-observation measurement error, s²_{ij} (Eq. S1). The linear predictor comprised the temperature function f(T_{ij}), the fixed bay-area and filtered-volume offsets (log A_j and log V_{ij}, the area term present only in the offset variants), and a bay-level random intercept α_j (Eq. S2–S3). Continuous predictors were z-standardised (mean centred, unit SD) before fitting; for the selected log-linear model the temperature term was z-scored log-temperature, so that f(T_{ij}) = β z_{logT,ij}, whereas the bay-area and filtered-volume offsets entered with their coefficients fixed at 1 and were not standardised. The measurement error was derived from the standard-curve maximum-likelihood fit as s_{ij} = σ_MLE / √n_reps. Adding this known error to the estimated residual variance, rather than relying on the residual alone, avoids underestimating uncertainty in individual Cq values, particularly at low concentrations. Full model implementation is provided in the supporting R script (Figshare: https://doi.org/10.6084/m9.figshare.32834141).

Eq. S1	$y_{ij}\sim\mathcal{N}\left( \mu_{ij}, \sigma^{2}+s_{ij}^{2} \right)$

Eq S2	$\mu_{ij}=\alpha_{j}+f\left( T_{ij} \right)+\log A_{j}+\log V_{ij}$

Eq S3	$\alpha_{j}\sim\mathcal{N}\left( \alpha,\sigma_{\alpha}^{2} \right)$

S1.3 Stage 1 priors (main analysis and prior sensitivity variants)

Because all continuous predictors were z-standardised (S1.1), priors were specified on the standardised scale, where a regression coefficient expresses the change in the eDNA index per standard deviation of the predictor; this makes a coefficient of large absolute value implausible a priori and allows weakly informative priors to be set on a common, interpretable scale (Gelman et al. 2008, Lemoine 2019). Priors common to all Stage 1 models were: intercept ∼ Normal(0, 5), placing the mean eDNA index within a broad but finite range; residual SD σ ∼ Exponential(1) and bay-level SD σ_α ∼ Exponential(1), weakly informative, strictly positive priors that regularise variance components toward zero without excluding plausibly large values, following the brms/Stan defaults (Carpenter et al. 2017, Bürkner 2017).

The temperature slope was the parameter of inferential interest, so we examined its sensitivity to three prior specifications spanning a deliberate range of informativeness. The weak prior, Normal(0, 2), is effectively uninformative on the standardised scale, admitting per-SD effects far larger than any plausible metabolic response and letting the data dominate (Gelman et al. 2017). The MMR-informed prior, Normal(0.44, 0.50), centres the slope on the metabolic expectation derived from the laboratory MMR Q10 of 3.5 (Cominassi et al. 2022), while remaining wide enough to be overridden by the data. The sceptical prior, Normal(0, 0.5), pulls the slope tightly toward zero and provides the most conservative test of whether a positive temperature effect persists under strong regularisation. Across these three priors, the back-transformed temperature slope ranged only from b = 4.28 (weak) to b = 3.82 (sceptical), corresponding to eDNA Q10 values that remained far above the metabolic benchmark in every case (Figure S 1).

S1.4 Bay area calculation and offset

Each bay was sampled with ten light traps (five unmodified, five modified). Only the five unmodified traps per bay contributed catch data to the analyses; all ten station positions were used to define the bay area (Figure S 2).

Bay areas were delineated from Lantmäteriet waterway shapefiles in the SWEREF 99 TM projection (EPSG:3006). For each bay, the convex hull of all ten station positions was buffered by a bay-specific radius (60 m for Bay C, 150 m for Bays A, B, and D) to encompass the spatial extent over which eDNA is diluted. The resulting areas were: Bay A = 0.041 km², Bay B = 0.027 km², Bay C = 0.063 km², Bay D = 0.018 km² (). The log-transformed area, log A_j, entered the eDNA models as a fixed offset (coefficient fixed at 1), accounting for spatial dilution of shed eDNA into larger or smaller water volumes.

S2 Quality control assessment of eDNA blanks and contamination risks
   S2.1 QC framework

Quality control of the qPCR data was conducted across three dimensions: (1) assessment of three distinct negative control types spanning the full workflow from field filtration to PCR amplification; (2) per-reaction inhibition assessment using an internal positive control (IPC) co-amplified in duplex with two *Esox lucius* assays on the Cy5 channel, run on the same extracted DNA as the *G. aculeatus* assay but in separate wells; and (3) evaluation of flagged field samples against a contamination threshold informed by the worst-case amplified negative control.

S2.2 Negative controls

Three negative control types were evaluated separately: field blanks (1 L MilliQ filtered in the field; detects environmental contamination during filtration), extraction blanks (detects reagent or bench contamination during DNA isolation), and PCR blanks (no-template controls; detects PCR setup contamination).

Field blanks amplified in 10 of 44 controls (22.7%; mean Cq 39.30 ± 2.17, range 35.1–43.3). All 10 detections occurred between 11 May and 21 June, coinciding with peak inshore *G. aculeatus* spawning activity (Bergström et al. 2015); no April blank amplified on any sampling date. To assess whether contamination inflated field measurements on dates when blanks did amplify, we compared the worst-case blank signal on each such date against the contemporaneous field sample concentrations. The strongest blank detected during the May–June period (Cq 35.1) corresponded to 0.91% of the mean field signal on those dates, indicating a negligible contamination contribution to contemporaneous field samples. Extraction blanks showed a single detection at Cq 42.63, at the extreme upper detection limit (~1–2 copies per reaction). PCR blanks were completely clean across all 12 replicates.

No field samples were excluded on contamination grounds. On dates when field blanks amplified (May–June), field signals were strong and the proportional contamination contribution was negligible. On April dates, all field blanks were non-detects, confirming that contamination was absent during filtration when field signals were weakest (Figure S 4).

S2.3 Internal positive control (IPC)

The IPC was detected in all reactions, with a mean Cq of 28.12 ± 0.43 SD and a total range of 27.2–29.6 across all sampling dates. No sample showed evidence of elevated IPC Cq relative to the plate mean, and no individual reaction was flagged for potential inhibition. These results indicate that PCR inhibition was absent at the level of the DNA extract across all samples. As the IPC was run in separate wells from the *G. aculeatus* assay, inhibition specific to the FAM channel cannot be fully excluded; however, such channel-specific inhibition is uncommon in environmental DNA applications and the absence of extract-level inhibition provides strong evidence that measurement error in the *G. aculeatus* assay reflects genuine qPCR variability rather than inhibition (Figure S 5).

S3 Supplementary figures

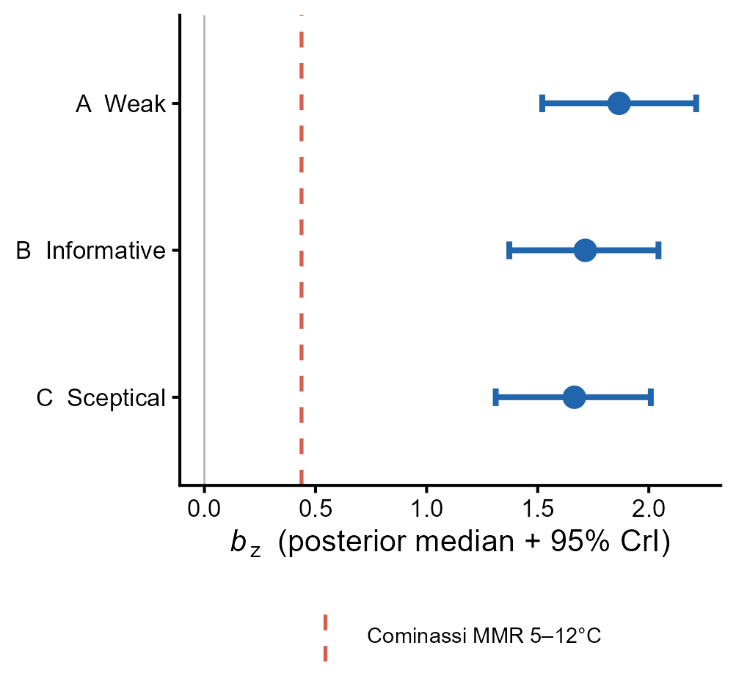

Figure S 1. Posterior estimates of the Stage 1 temperature slope (b_z, standardised scale) under three prior specifications: weak (A), MMR-informed (B), and sceptical (C). The dashed red line indicates the MMR-equivalent slope. All three posteriors yield Q10 values substantially exceeding the MMR benchmark (Q10 = 3.5).

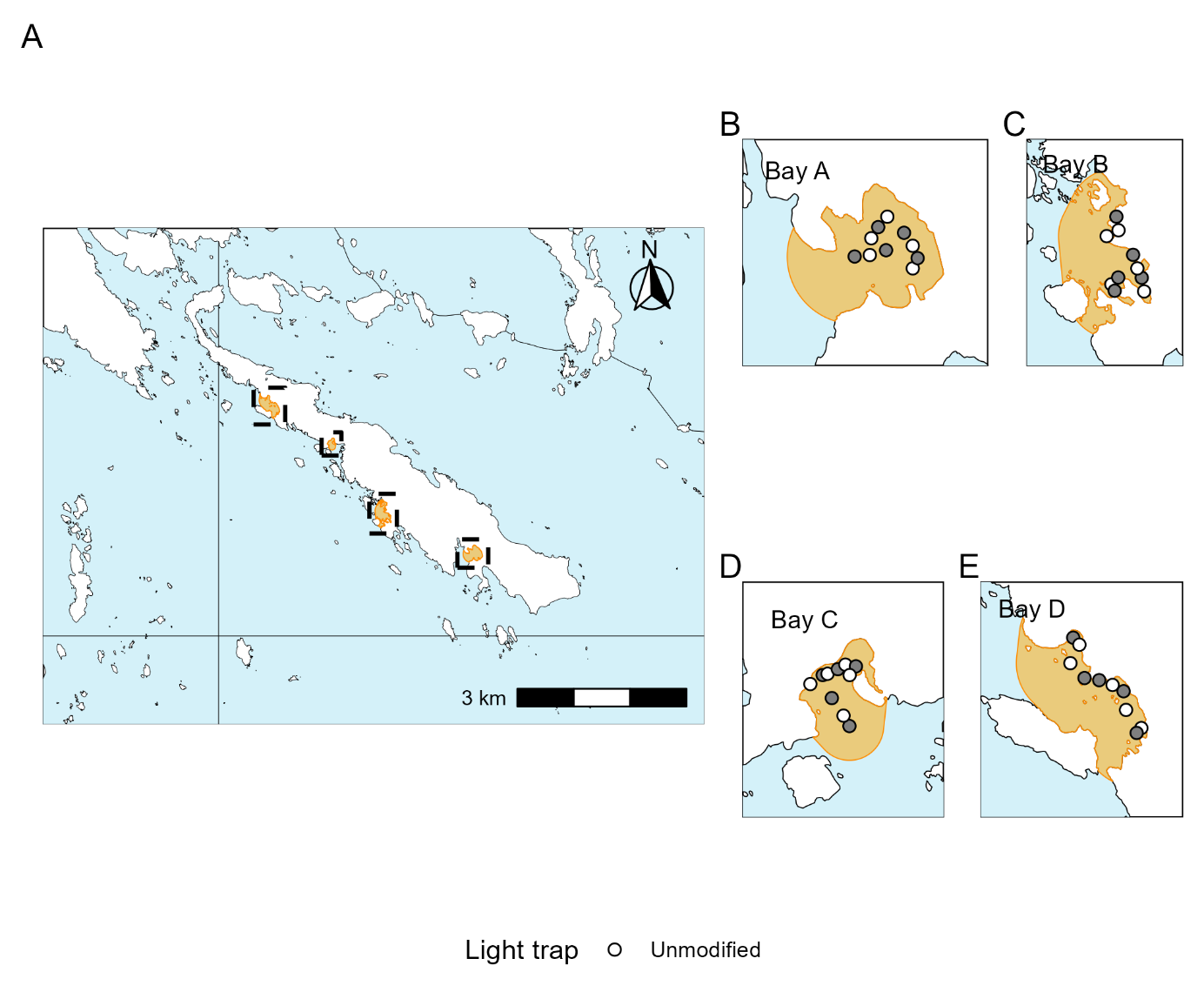

Figure S 2. Spatial dilution basins at the four study bays (A, regional overview; B–E, per-bay detail). Points show station positions where both light traps and benthic traps were moored. Two types of light traps were deployed (Ogonowski 2026) but only catches from the “Unmodified” traps were used in all analyses. All mooring stations within bays contributed to the the calculation of bay area. Orange polygons show the convex hull of all ten station positions, buffered (bay-specific radius 60–150 m) to encompass the sampled water area; this area defines the bay-level dilution offset (log A_j).

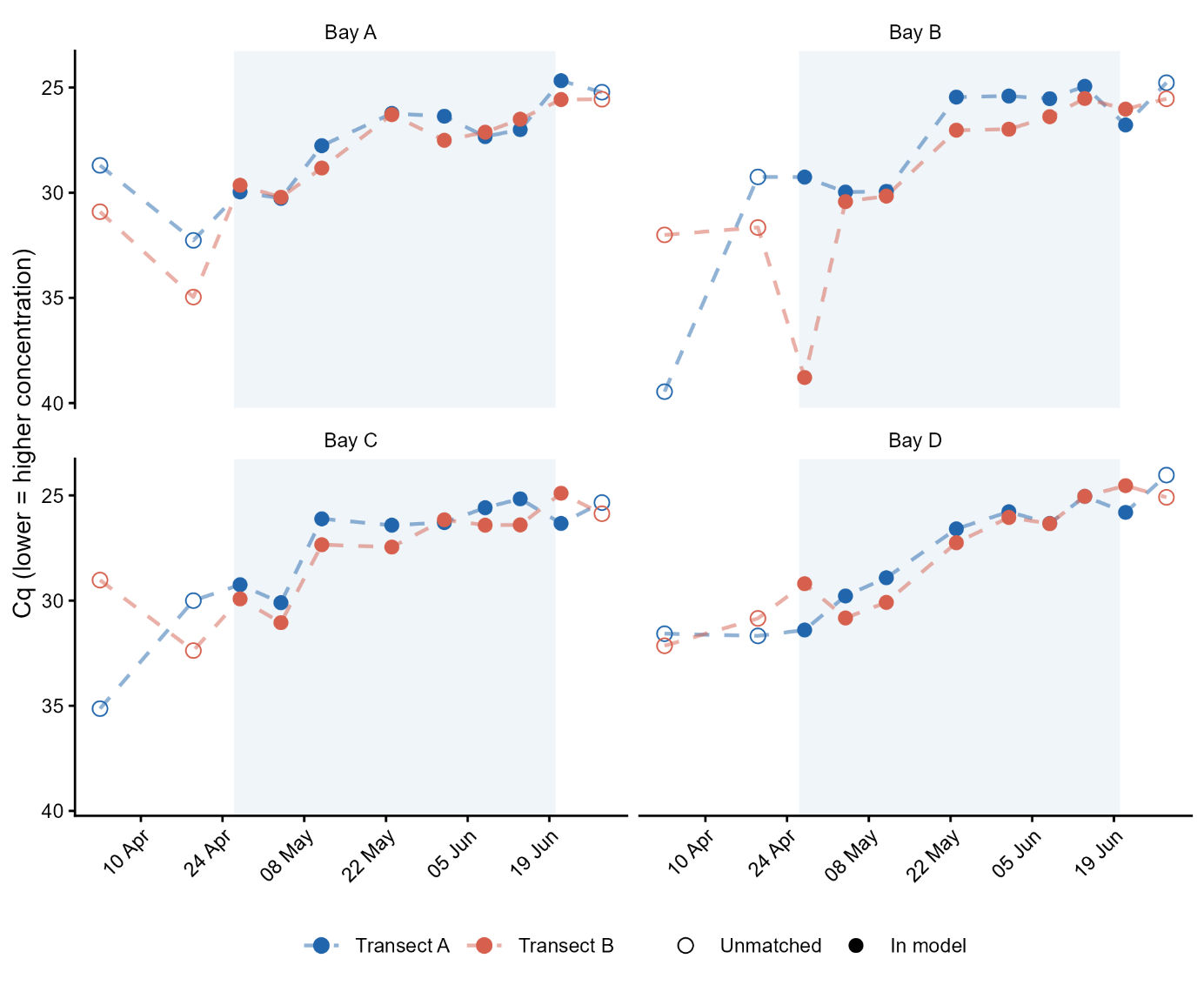

Figure S 3. Full seasonal trajectory of eDNA Cq values per bay × transect, including post-trap-deployment unmatched observations used to estimate in situ degradation. Filled circles indicate samples included in the main analysis (in model); open circles indicate unmatched observations outside the matched analysis window. The shaded blue rectangle indicates the matched dataset window used in the main analysis.

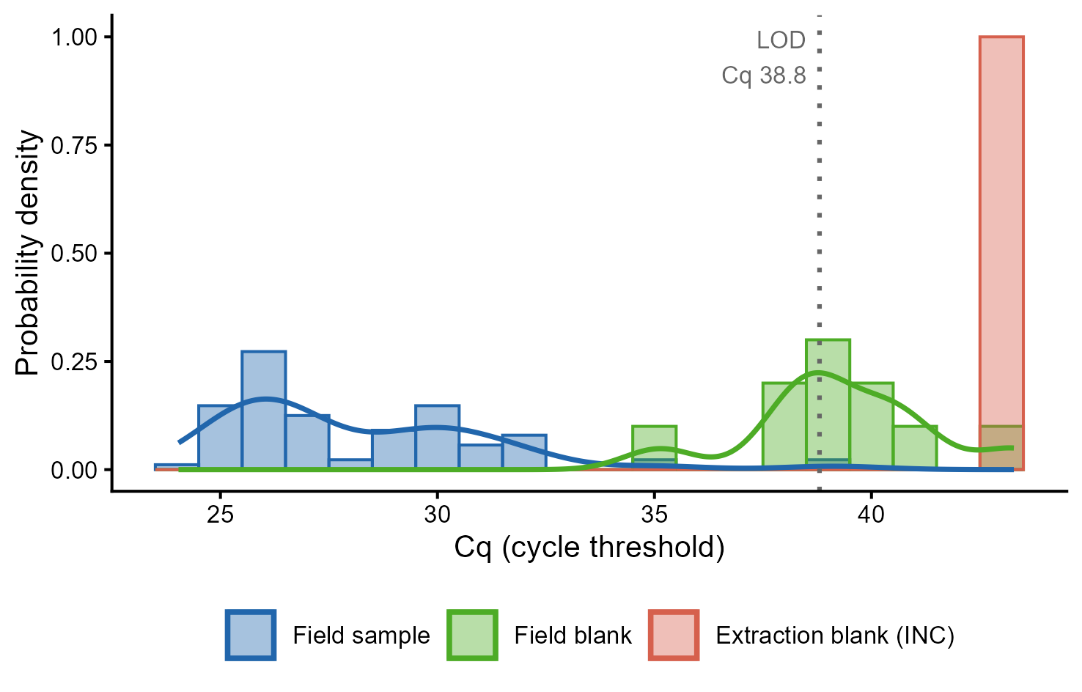

Figure S 4. Distribution of Cq values for field samples and the two detected negative control types (field blanks and extraction blanks). The dotted grey line indicates the limit of detection (LOD, Cq 38.80). PCR blanks are excluded from the figure as all 12 were non-detects.

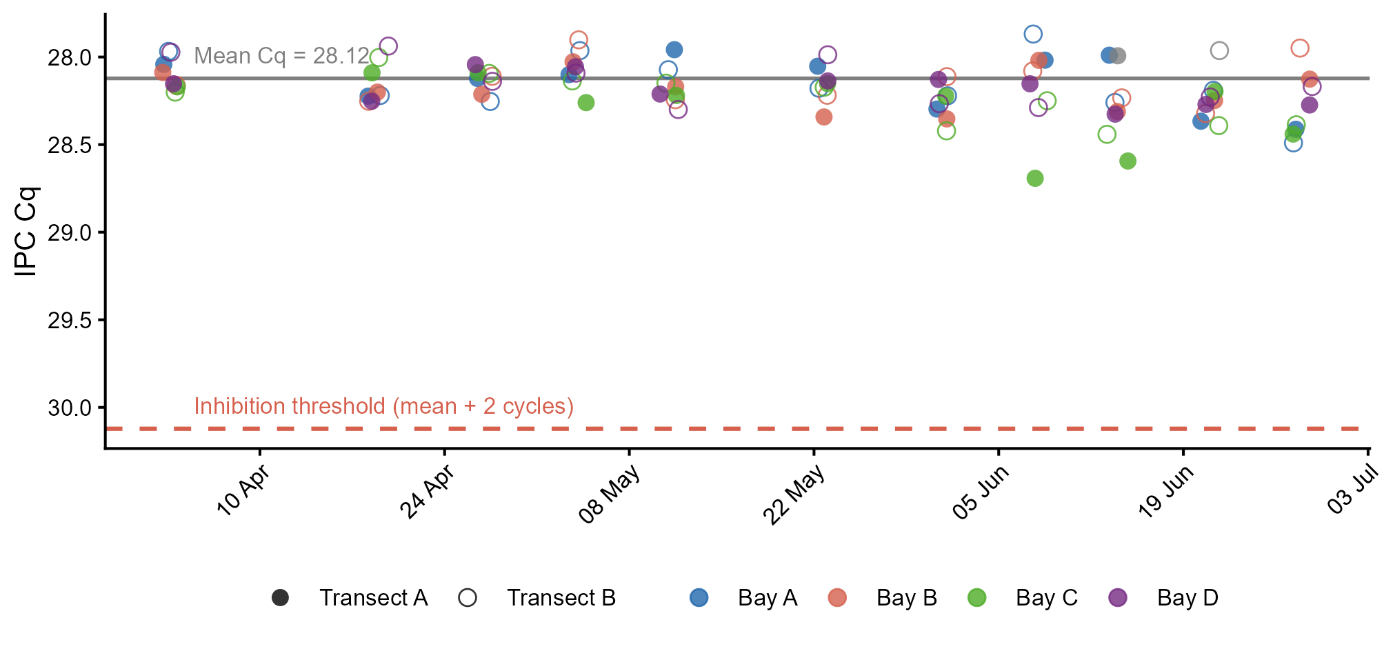

Figure S 5. Internal positive control (IPC) Cq per field sample, coloured by bay and shaped by transect. The IPC was co-amplified in the same reaction well as the G. aculeatus FAM assay on the Cy5 channel. The horizontal solid grey line indicates the overall mean IPC Cq (28.12); the dashed red line indicates the inhibition threshold (mean + 2 cycles = 30.12). No sample was flagged for potential inhibition.

S4 Supplementary tables

Table S 1. Leave-one-out (LOO) cross-validation comparison (Vehtari et al. 2017) of six candidate models for the eDNAindex ~ Temperature relationship. Response variable: edna_index = −mean Cq with brms se (se_edna, sigma = TRUE) measurement error propagation. Three functional forms (linear, log-linear/Arrhenius, quadratic) were crossed with two bay area-offset decisions (“na” denotes no area offset). **elpd_diff**: difference in elpd_loo from the top-ranked model. **se_diff**: standard error of elpd_diff. **p_loo**: estimated effective number of parameters.

| **Model** | **elpd_diff** | **se_diff** | **p_loo** |
| --- | --- | --- | --- |
| quad_na | 0.00 | 0.00 | 13.00 |
| quadratic | -0.81 | 1.01 | 14.07 |
| log_na | -1.55 | 1.85 | 13.73 |
| log | -2.11 | 1.26 | 14.22 |
| linear_na | -4.45 | 2.00 | 11.95 |
| linear | -4.94 | 2.98 | 12.34 |
